# Bayesian phylogenetics on globally emerging SARS-CoV-2 variant BA.2.86 suggest global distribution and rapid evolution

**DOI:** 10.1101/2023.09.08.556912

**Authors:** Andrew P. Rothstein, Xueting Qiu, Keith Robison, Susan Collins, Gabi Muir, Bernadette Lu, Alex M. Plocik, Birgitte B. Simen, Casandra W. Philipson

## Abstract

Using bioinformatic pipelines and Bayseian phylogenetic analyses, we characterized a SARS-CoV-2 variant designated by the World Health Organization as a variant under monitoring in August 2023. Here we analyze the genomes of this SARS-CoV-2 variant, BA.2.86, deposited into GISAID within the two weeks of its emergence (2023-08-14 first submission to 2023-08-31), including the first BA.2.86 genome reported from a traveler originating from Japan. We present bioinformatics methods using publicly available tools to help analysts identify the lineage-defining 12 nucleotide insertion (S:Ins16MPLF), which is often masked by most bioinformatics pipelines. We also applied maximum-likelihood and Bayesian phylogenetics to demonstrate the high mutational rate of the tree branch leading to the emergence of BA.2.86, hinting at possible origins, and predict that BA.2.86 emerged around May 2023 and spread globally rapidly. Taken together, these results provide a framework for more rigorous bioinformatics approaches for teams performing genomic surveillance on viral respiratory pathogens.

## Introduction

On August 13th, 2023, Israel published the first genome of a divergent severe acute respiratory syndrome coronavirus 2 (SARS-CoV-2) variant, designated BA.2.86, with a notable number of S-gene mutations (>30 amino-acid substitutions) in the Spike protein compared to BA.2^1^. Subsequently, BA.2.86 was detected in multiple countries including Denmark, the United Kingdom, South Africa, and the United States. Based on the global detection and mutational profile, on August 17th the World Health Organization (WHO) quickly classified BA.2.86 as a variant under monitoring^2^. Given the unique profile of BA.2.86, a number of standard bioinformatic pipelines have had difficulty identifying key BA.2.86 defining mutations, specifically a lineage-defining 12 nucleotide insertion. This region is unresolved in many of the earliest publicly deposited genomes, which made it difficult to infer within BA.2.86 phylogenetic relationships. Due to the rapid emergence and importance of BA.2.86, we sought to compare available BA.2.86 sequence data from public repositories as of 2023-08-31 to glean insights on bioinformatic challenges and evolution of the virus. In this study we compare available BA.2.86 consensus genomes, strategize for adjusting bioinformatic pipelines to capture lineage-defining mutations for BA.2.86, and, given public repository submissions may be a poor representation of true variant detection timing, apply Bayesian phylogenetic approaches to infer the timing of BA.2.86 emergence. Our Bayseian phylogenetic results demonstrate a high mutational rate of the tree branch leading to the emergence of BA.2.86, hinting at possible origins, and predict that BA.2.86 emerged around May 2023 and spread globally rapidly.

## Methods

### Data

We analyzed all BA.2.86 genomes deposited into GISAID within the two weeks of the variant’s emergence (2023-08-14 first submission to 2023-08-31). Consensus genomes were retrieved from GISAID. Specifically, EPI_ISL_18097345, EPI_ISL_18125259, EPI_ISL_18125249, EPI_ISL_18138566, EPI_ISL_18096761, EPI_ISL_18097315, EPI_ISL_18164467, EPI_ISL_18110065, EPI_ISL_18114953, EPI_ISL_18159709, EPI_ISL_18160063, EPI_ISL_18147545, EPI_ISL_18121060, EPI_ISL_18111770, EPI_ISL_18151536, EPI_ISL_18135401, EPI_ISL_18158884, EPI_ISL_18157710, EPI_ISL_18158448, EPI_ISL_18142273, EPI_ISL_18142275, EPI_ISL_18168185, EPI_ISL_18147561, EPI_ISL_18168405, EPI_ISL_18147559, EPI_ISL_18158452, EPI_ISL_18168588, EPI_ISL_18164159. Raw sequencing data for a genome from a traveler arriving from Japan on August 10th, 2023^3^ (GISAID EPI_ISL_18121060) was also made available in NCBI’s Sequencing Read Archive (SRR25750759). Findings of this study are based on sequences and metadata associated with the 28 BA.2.86 sequences available on GISAID up to August 31, 2023 and accessible at doi:10.55876/gis8.230831ur. Global lineages for the 83 sequences used in the Bayesian phylogenetic analysis are accessible at doi:10.55876/gis8.230902wk.

### Bioinformatics analysis

Four custom in-house pipelines were tested to generate a consensus sequence, assign the BA.2.86 lineage, and confirm the lineage-defining 12nt insertion sequence, GTGTGTCATGCCGCTGTTTAAT. The four pipelines vary primarily in variant calling and assembly methods. Specifically, Pipeline 1 implemented the most strict variant calling parameter settings to identify SNPs and implemented soft clipping of adapter sequences to minimize bioinformatics artifacts; Pipeline 2 implemented more relaxed variant calling parameter settings and omitted soft clipping; Pipeline 3 substituted a variant caller known to perform well on small (<50nt) indels, DeepVariant; Pipeline 4 performed *de novo* assembly in lieu of reference-based assembly. Detailed tools and parameter settings can be found in the Supplementary Information.

### Phylogenetics and mutational profiles

Consensus genome sequences (n=30) were aligned and mutational profiles were generated using Nextclade (v2.14.1)^4^. Consensus reference genomes for XBB.1.5 and BA.2 were accessed here: https://github.com/corneliusroemer/ncov-simplest/tree/main/data. A maximum likelihood phylogenetic tree was generated using the aligned sequences (iqTREE v1.6.12, 1,000 bootstraps)^5^. iqTREE model finder was implemented and HKY+F+I was chosen according to Bayesian Information Criterion. The maximum likelihood tree was visualized and annotated using iTOL (v6)^6^.

### Ancestral sequence reconstruction

The ancestral sequence of BA.2.86 was reconstructed using TreeTime. TreeTime enables reconstructing likely sequences of internal nodes of the phylogenetic tree and modeling how sequences change along the tree^7^. The ancestral sequence was inferred by the general time reversible (GTR) nucleotide substitution model. To input the high quality sequences of BA.2.86, out of the 28 sequences, three sequences (EPI_ISL_18147559, EPI_ISL_18147561, and EPI_ISL_18160063) were removed based on the Nextclade’s reporting of frame shifts. The final 25 sequences (Supplementary Information 1) of BA.2.86 were aligned with Muscle v3.8.31^8^ and manually verified to ensure the quality of the data for ancestral sequence reconstruction, thus, the alignment is not aligned to a reference genome. The reconstructed ancestral sequence and mutations on the tree branches were reported. The phylogenetic tree was midpoint rooted.

### Bayesian Phylogenetics

Bayesian phylogenetic analysis for BA.2.86 was conducted on 2023-08-31, including the referenced BA.2.86 sequences (n=28). A random SARS-CoV-2 sequence dataset (n=83) was generated by repeating maximum-likelihood phylogenetic analysis from GISAID to represent the global historical and circulating lineages, bringing the final dataset to a total of 111 sequences. All sequences were aligned with Muscle v3.8.31^8^ and manually verified to ensure the quality of the data. The Bayesian phylogenetic analysis was performed in a Bayesian statistical framework implemented in BEAST (v.1.10.4)^9^. A general time reversible (GTR) nucleotide substitution model was used with a 4-category gamma distribution of variation among sites and a proportion of invariant sites. A log-normal uncorrelated relaxed molecular clock was applied with an initial mean of 0.0008 and a uniform prior ranging from 0.0 to 1.0. The Bayesian Skyride coalescent model estimating the effective population size was used with a Gaussian Markov random field (GMRF) smoothing prior on the parameters of the piecewise constant population size trajectory^10^. Two independent Markov chain Monte Carlo (MCMC) simulations of 100-million steps converged and provided Bayesian estimates on the emerging ancestral time and evolutionary rate of BA.2.86. To ensure proper convergence and parameter mixing with an effective sample size (ESS) of at least 200, a minimum of 10% burn-in was removed, that is, samples at the beginning of the MCMC run prior to its convergence to the stationary distribution were discarded. TreeAnnotator version 1.10.4^9^ was used to generate the maximum clade credibility (MCC) phylogenetic tree, which was visualized in FigTree v1.4.2^11^. The most recent common ancestor (TMRCA) and 95% BCI (Bayesian credible interval) was estimated for the BA.2.86.

## Results

### BA.2.86 genomic description

We analyzed all BA.2.86 genomes deposited into GISAID within the two weeks of the variant’s emergence (2023-08-14 first submission to 2023-08-31). This first set of sequences were submitted by labs across five continents including countries of Denmark (n=10), Sweden (n=5), South Africa (n=3), United States (n=4, including 1 sample collected from a traveler returning from Japan to the US), Portugal (n=2), Canada (n=1), France (n=1), Israel (n=1), and the United Kingdom (n=1). The average reference coverage of the publicly available consensus genomes was 97.6%. Compared to the reference BA.2, BA.2.86 genomes have an average of 51.5 additional substitutions and 53.3 deletions (**Table 1**). On average each BA.2.86 sequence has the presence of 4.9 private mutations relative to BA.2. Notably, the BA.2.86 branch included at least 27 divergent mutations within SARS-CoV-2 S-gene compared to BA.2 (including R21T, S50L,H69-, V70-, V127F, Y144-, F157S, R158G, N211-, L212I, L216F, H245N, A264D, I332V, K356T, R403K, V445H, N450D, L452W, N481K, V483-, E484K, F486P, E554K, A570V, P621S, P681R, S939F, P1143L).

**Table 1.** Summary mutation statistics for BA.2.86 samples from Nextclade. Total mutational counts are relative to SARS-CoV-2 reference. Immune Escape and ACE2 binding scores are relative to BA.2.

| Sequence Name | GISAID ID | Date Collected | Pangolin Lineage | Immune Escape | ACE2 Binding | Total Substitutions | Total Deletions | Total Insertions | Total Private Mutations |
| --- | --- | --- | --- | --- | --- | --- | --- | --- | --- |
| hCoV-19/Denmark/DCGC-647676/2023 | EPI_ISL_18097345 | 2023-07-24 | BA.2.86 | 1.20 | 1.05 | 49 | 30 | 0 | 0 |
| hCoV-19/South_Africa/NICD-N55967/2023 | EPI_ISL_18125259 | 2023-07-24 | BA.2.86 | 0.84 | 1.29 | 46 | 65 | 0 | 6 |
| hCoV-19/South_Africa/NICD-N55999/2023 | EPI_ISL_18125249 | 2023-07-28 | BA.2.86 | 1.03 | 1.16 | 54 | 65 | 0 | 10 |
| hCoV-19/USA/OH-ODH-SC3032044/2023 | EPI_ISL_18138566 | 2023-07-29 | BA.2.86 | 1.20 | 1.05 | 51 | 68 | 12 | 3 |
| hCoV-19/Israel/ICH-741198454/2023 | EPI_ISL_18096761 | 2023-07-31 | BA.2.86 | 1.20 | 0.95 | 54 | 68 | 12 | 0 |
| hCoV-19/Denmark/DCGC-647646/2023 | EPI_ISL_18097315 | 2023-07-31 | BA.2.86 | 1.20 | 1.05 | 51 | 21 | 0 | 0 |
| hCoV-19/Canada/BC-BCCDC-641728/2023 | EPI_ISL_18164467 | 2023-08 | BA.2.86 | 1.20 | 1.05 | 51 | 42 | 12 | 5 |
| hCoV-19/USA/MI-UM-10052670540/2023 | EPI_ISL_18110065 | 2023-08-03 | BA.2.86 | 1.20 | 1.05 | 53 | 42 | 12 | 2 |
| hCoV-19/Denmark/DCGC-647694/2023 | EPI_ISL_18114953 | 2023-08-07 | BA.2.86 | 1.20 | 1.05 | 52 | 42 | 12 | 4 |
| hCoV-19/Denmark/DCGC-658505/2023 | EPI_ISL_18159709 | 2023-08-07 | BA.2.86 | 1.20 | 1.05 | 52 | 42 | 12 | 4 |
| hCoV-19/Denmark/DCGC-658800/2023 | EPI_ISL_18160063 | 2023-08-07 | BA.2.86 | 0.00 | 0.00 | 49 | 43 | 12 | 6 |
| hCoV-19/Sweden/AB-000-117-986-1/2023 | EPI_ISL_18147545 | 2023-08-07 | BA.2.86 | 0.54 | 0.44 | 40 | 53 | 0 | 2 |
| hCoV-19/USA/VA-GBW-H20-330-6734/2023 | EPI_ISL_18121060 | 2023-08-10 | BA.2.86 | 1.20 | 1.05 | 53 | 68 | 12 | 8 |
| hCoV-19/England/GSTT-YYBYBN4/2023 | EPI_ISL_18111770 | 2023-08-13 | BA.2.86 | 1.20 | 1.05 | 53 | 68 | 12 | 5 |
| hCoV-19/Sweden/AB-01_SE100_CS101480/2023 | EPI_ISL_18151536 | 2023-08-13 | BA.2.86 | 0.89 | 1.05 | 53 | 68 | 12 | 9 |
| hCoV-19/Denmark/DCGC-647853/2023 | EPI_ISL_18135401 | 2023-08-14 | BA.2.86 | 1.20 | 1.05 | 53 | 30 | 0 | 4 |
| hCoV-19/Denmark/DCGC-657658/2023 | EPI_ISL_18158884 | 2023-08-14 | BA.2.86 | 1.20 | 1.05 | 53 | 42 | 12 | 5 |
| hCoV-19/Denmark/DCGC-656489/2023 | EPI_ISL_18157710 | 2023-08-14 | BA.2.86 | 1.20 | 1.05 | 52 | 42 | 12 | 5 |
| hCoV-19/Denmark/DCGC-657228/2023 | EPI_ISL_18158448 | 2023-08-14 | BA.2.86 | 1.20 | 1.05 | 52 | 42 | 12 | 5 |
| hCoV-19/Portugal/PT49343/2023 | EPI_ISL_18142273 | 2023-08-15 | BA.2.86 | 1.19 | 0.32 | 52 | 68 | 0 | 6 |
| hCoV-19/Portugal/PT49347/2023 | EPI_ISL_18142275 | 2023-08-15 | BA.2.86 | 1.19 | 0.32 | 52 | 68 | 0 | 6 |
| hCoV-19/South_Africa/NICD-R09343/2023 | EPI_ISL_18168185 | 2023-08-17 | BA.2.86 | 0.89 | 1.49 | 48 | 65 | 0 | 7 |
| hCoV-19/Sweden/Y-261733286655/2023 | EPI_ISL_18147561 | 2023-08-18 | BA.2.86 | 0.72 | 0.20 | 53 | 79 | 0 | 7 |
| hCoV-19/Sweden/U-M988749/2023 | EPI_ISL_18168405 | 2023-08-18 | BA.2.86 | 1.20 | 1.05 | 53 | 42 | 12 | 5 |
| hCoV-19/Sweden/Y-261733291192/2023 | EPI_ISL_18147559 | 2023-08-19 | BA.2.86 | 0.72 | 0.20 | 52 | 52 | 0 | 6 |
| hCoV-19/Denmark/DCGC-657232/2023 | EPI_ISL_18158452 | 2023-08-21 | BA.2.86 | 1.20 | 1.05 | 53 | 42 | 12 | 5 |
| hCoV-19/France/GES-IPP17379/2023 | EPI_ISL_18168588 | 2023-08-21 | BA.2.86 | 1.20 | 1.05 | 53 | 68 | 12 | 5 |
| hCoV-19/USA/TX-HMH-M-132251/2023 | EPI_ISL_18164159 | 2023-08-22 | BA.2.86 | 1.20 | 1.05 | 55 | 68 | 12 | 7 |
| XBB.1.5 (reference) | N/A | N/A | XBB.1.5 | 0.91 | 1.01 | 29 | 56 | 0 | 1 |
| BA.2 (reference) | N/A | N/A | BA.2 | 0.00 | 0.00 | 0 | 53 | 0 | 0 |

BA.2.86 sequences contain a number of S-gene mutations predicted to confer greater antibody escape and ACE2 binding affinity. Based on deep mutational scanning approaches^12,13^, the BA.2.86 S-gene mutations that are predicted to increase antibody escape relative to BA.2 are ins16MPLF, Y144-, F157S, R158G, H245N, A264D, K356T, V445H, G446S, N450D, L452W, N460K, -V483, and A484K. Notably, in vitro studies have highlighted that S-gene mutations N450D, K356T, L452W, A484K, V483-, V445H compared to XBB.1.5 were most important for neutralizing antibodies in pseudovirus experiments^14^. In addition, the S-gene mutations present in BA.2.86 that are predicted to increase ACE2 binding affinity relative to BA.2 are based on the mutations R403K, N460K, and R493Q. Despite the prediction of increased ACE2 binding affinity compared to BA.2, when compared to XBB.1.5 *in vitro*, the presence of K356T, V483del, and E554K lowered BA.2.86 infectivity^14^.

### Bioinformatic challenges for reference-based assembly of BA.2.86 genomes

One of the most defining characteristics of the BA.2.86 lineage is a 12 nucleotide insertion (S:Ins16MPLF; GTGTGTCATGCCGCTGTTTAAT) at the start of the S gene. Among the first 28 publicly available consensus genomes, 17 (60%) reported this insertion while the remaining 11 genomes (40%) were masked (e.g. Ns or “-”) across the region. This region could be missed for a variety of reasons including sequencing platform, amplicon based primer schemes that are designed adjacent to the insertion location (such as 70_LEFT in ARTIC primer schemes^15^), default parameters for bioinformatic assembly, as well as thresholds for variant detection. Extensive soft clipping of reads is often performed in order to avoid primer artifacts, especially for labs employing widely accepted amplicon-based short-read sequencing, however indels in the true sequence can lead to incorrect trimming of very short primer or adapter derived sequences. In other words, the evidence for calling indels strongly resembles the evidence for identifying common sequencing and alignment artifacts. For these reasons, we investigated possible root causes for bioinformatics artifacts observed in BA.2.86 genomes in public repositories.

### Masking of the 12 nucleotide insertion (S:Ins16MPLF) due to minimum alternate fraction parameter setting in freebayes

The 12 nucleotide insertion in the S gene should be called MN908947.3 21608 . GTTAA GTCATGCCGCTGTTTAA. In our experience, we find in our sample (SRR2575079) and others that whether freebayes 1.3.6 calls this is dependent on the setting of --min-alternate-fraction parameter. Setting this parameter to <=0.42 results in the insertion being identified in our dataset. However, if the parameter is above the 0.42 threshold, then the insertion is masked.

We also observe that the –min-alternate-fraction parameter is sensitive to reference coverage at T21609 when reads do not extend into the deletion, resulting in the reference mutation being called instead of reversion (G22200Trev). After performing additional testing, we believe this behavior is occurring due to an issue, which we have submitted to the freebayes team for consideration via GitHub (https://github.com/freebayes/freebayes/issues/775).

### Resolving consensus genomes using orthogonal bioinformatics pipelines

We ran 4 pipelines (Supplementary Information) on our data that varied primarily in variant calling and assembly methods to resolve the BA.2.86 genome (EPI_ISL_18121060). This sequence, identified in a traveler from Japan, is unique in its phylogenetic placement among BA.2.86 lineages (**Figure 1**). Compared to the other consensus sequences analyzed, this sample is defined by a distinct branch with four unique nucleotide mutations C222T, C1960T, T12775C, and G13427A as well as one amino acid substitution ORF1a:E4388L. Prioritizing high-frequency variant calls with strict freebayes parameters (Pipeline 1) resulted in masking of the S:Ins16MPLF insertion. However, using outputs from multiple pipelines with more relaxed variant calling filtering (Pipeline 2), or other variant callers including the neural network-based variant caller DeepVariant (Pipeline 3), or de novo assembly (Pipeline4) successfully identified the S:Ins16MPLF insertion. Results from all four pipelines were compared manually (visual inspection of read mapping and variant calling) and used to generate a polished consensus genome.

**Figure 1.**
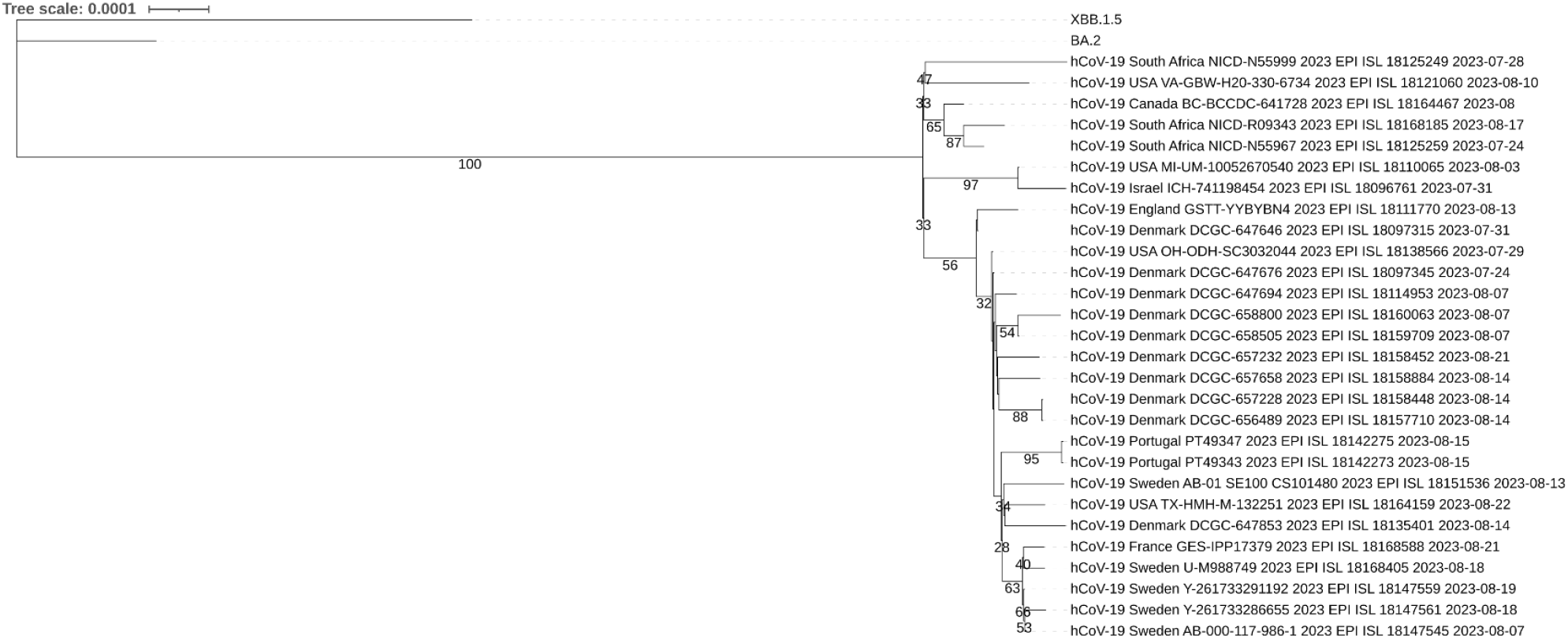
Maximum likelihood phylogenetic tree (model HKY+F+I) of all 28 public BA.2.86 genomes and reference groups of BA.2 and XBB.1.5. Branch labels indicate bootstrap support based on 1,000 bootstraps. BA.2.86 samples cluster as a well supported clade with some phylogenetic structuring. The sample from a traveler arriving in the USA from Japan is embedded within BA.2.86 but has support (47% bootstrap support) for distinct branching.

### Ancestral sequence reconstruction

The reconstructed ancestral sequence and the phylogenetic tree mapped with mutations on the branches can be found in the Supplementary data 1 and Figure S1. The USA-MI isolate (EPI_ISL_18110065) was inferred as the ancestral sequence and the Israel isolate (EPS_ISL_18096761) were two mutations away from the USA-MI isolate. Given the impacts of potential errors in the sequences, the clade of USA-MI and Israel isolates were the phylogenetically “oldest” sequences, indicating they are most close to the inferred ancestral sequence, out of the about three-week time window of sequences collected for BA.2.86.

### Bayesian Phylogenetic Analysis

The ancestral time of the BA.2.86 lineage divergence from other Omicron lineages was estimated as 2022.23 (95% HPD: 2022.05, 2022.44), which was rooted and similar to the original BA.2 emerging time (around March 26th, 2022) (Figure 2). The common ancestral time of the 28 BA.2.86 sequences was 2023.32 (95% HPD: 2023.21, 2023.42), which emerged and started to spread in the population around May 1st of 2023 (95% HPD: March 21, 2023; June 2, 2023). Our inference for the time to the most recent common ancestor is consistent with previous estimates^16^. The BA.2.86 branch evolutionary rate was estimated as 0.0013 (95% HPD: 0.0009, 0.0018) substitutions per site per year, on the higher rate compared to the average evolutionary rate of SARS-CoV-2 (about 0.0008 substitutions per site per year). Among the 28 BA.2.86 sequences, isolates from European countries (including Sweden, France, Denmark, Portugal and England) are more closely clustered; isolates from South Africa clustered with sequences from Canada the traveler from Japan to the USA; the Israel sample is clustered with the USA-MI sample, indicating that BA.2.86 has been widely spreading globally. More data is needed to infer the spatial connections among the geographic locations.

**Figure 2.**
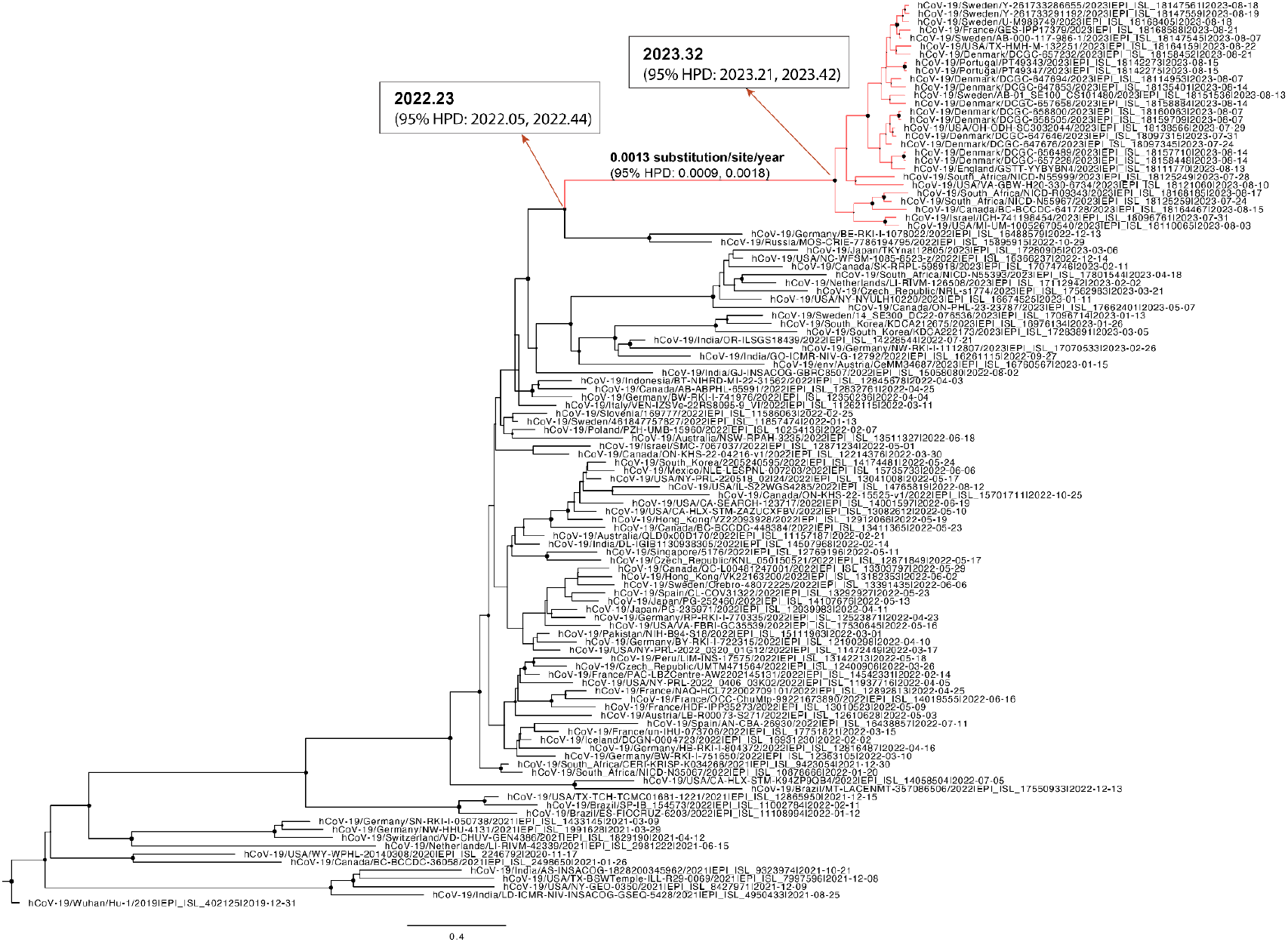
BEAST phylogenetic tree. Red branches are those of BA.2.86. Branch lengths indicate genetic distances on the year time scale. Size of node circles indicate posterior probability ranging from 0-1, where the value of >= 0.7 indicates statistical significance on the divergence. Black dots on the tree nodes indicate BA.2.86 posterior probabilities within clade. Red arrows and corresponding box highlight ancestral timing for BA.2.86 diverging from the BA.2 root and its emergence/spread in the population.

In summary, the Bayesian phylogenetic analysis indicates that the evolutionary rate leading to the emergence of BA.2.86 was faster compared to the rate in the general population. Given the evolutionary rate is above baseline SARS-CoV-2, a central hypothesis is that evolution and divergence could be happening either in immunocompromised populations before widespread, or outside human populations (alternative hosts, independent evolution). These are similar hypotheses to the original Omicron’s origin and emergence.

## Discussion/Conclusions

### BA.2.86 has important mutational differences compared to known circulating SARS-CoV-2 variants

Despite BA.2 and descendants’ dominance during Mar-June 2022, BA.2 was replaced by waves of BA.4/BA.5 in late 2022 followed by the emergence and global spread of XBB for the majority of 2023^17^. By December 2022, XBB, specifically XBB.1.5, was the dominant globally circulating lineage^2^ and on May 18th, 2023 was recommended by the WHO as the variant choice for 2023-2024 SARS-CoV-2 global vaccine booster^18^. Therefore, it is important to highlight mutations present in BA.2.86 that may confer an antigenic advantage relative to past variant exposure histories among the global human population.

In comparison to XBB.1.5^19^, there are at least 28 SARS-CoV-2 spike mutations only present in BA.2.86, at least eleven mutations only present in XBB.1.5, and 31 shared mutations. Mutations present in BA.2.86 at sites (e.g. S:R356T, S:L452W) are absent in XBB.1.5 and are predicted to increase antibody escape. In addition, further mutations such as S:V445H (compared to S:V445P in XBB.1.5) and S:E484K (compared to S:E484A in XBB.1.5) are expected to add additional immune escape on top of already selectively advantageous sites with SARS-CoV-2 spike protein^12^. However, even with a large number of distinct and shared mutations, it is currently unknown how well BA.2.86 will outcompete dominant traits of XBB and its descendants.

*In vitro* data to date has indicated that BA.2.86 is antigenically distinct from other previous variants based on cartography and has strong immune evasive properties based on challenges against against XBB* infection or vaccine induced antibodies^20–22^. However, relative to XBB.1.5, BA.2.86 exhibited lower pseudovirus infectivity suggesting this may impact transmission efficiency^20^. Despite lower inferred transmissibility relative to other dominant circulating lineages, more data is needed on the diversity of BA.2.86. It is still not clear whether undetected BA.2.86 lineages have known mutations that could increase infectivity (e.g. ACE2 binding mutations). If BA.2.86 were to acquire known infectivity mutations (such as combinations of S:L455F & S:F456L or S:Q498R & N501Y which is epistatic effect have historically increased ACE2 binding among SARS-CoV-2 variants)^23,24^ along with the current repertoire of mutations, there may be an opportunity for BA.2.86 to quickly outcompete globally dominant variants.

### Inferring origin of BA.2.86 requires investigations in immunocompromised and animal host populations

The origin of the original Omicron lineage that emerged at the end of 2021 is still unknown and the pattern and profile of BA.2.86 seems to repeat this phenomenon of emergence with a large set of unique mutations in mid-2023. From the phylogenetic structure and a long branch leading to the emergence of BA.2.86, it resembles the emergence of the initial Omicron lineage. Three potential theories have been proposed to explain the origin of Omicron: 1) Omicron may have circulated and evolved in an unsampled population; 2) Omicron may have evolved from a host-pathogen evolutionary race between immunosuppressed patients and SARS-CoV-2; 3) Omicron could have originated from adaptation in animal reservoirs and been transmitted back to humans (i.e. zoonotic spillback)^25^. Intense global genomic surveillance for SARS-CoV-2 during 2021-2022 generally refutes the possibility of the first theory (i.e., circulating in a hidden population), leaving immunocompromised patients or animal reservoir hosts as plausible hypotheses.

Bayesian phylogenetic analysis in this study estimated a high branch evolution rate that led to the emergence of BA.2.86, indicating that since its divergence from BA.2 in early 2022, over the past year, BA.2.86 has been evolving in a host or a host population before its dissemination in the human population recently. Both the immunocompromised host theory and animal host theory can be supported by this rapid evolutionary rate of BA.2.86. A meta-analysis study of SARS-CoV-2 persistent infection individuals showed that SARS-CoV-2 evolution was significantly faster in these persistent infected immunocompromised patients than that in the general human population^26^. A recent cohort study also reported that although prolonged replication-competent Omicron SARS-CoV-2 infections were uncommon, the within-host evolution reported an increased nonsynonymous mutation rate in persistent patients compared to short infections, where the persistently infected individuals accumulated globally distinct Spike mutations^27^. Animal spillover and human-to-animal SARS-CoV-2 spillback have been frequently reported^28^. For example, the Omicron lineage origin could be the result of SARS-CoV-2 transmitting back from mice after adaptation. Several studies have demonstrated Omicron mutations generated in mouse-adapted SARS-CoV-2 variants by serial passaging in mice^29^. A recent study on SARS-CoV-2 in white-tailed deer in Ohio, USA during November 2021 to March 2022, showed that the SARS-CoV-2 virus evolved not only three-times faster in white-tailed deer compared to the rate in humans, but was also driven by different mutational biases and selection pressure^30^. Therefore, both in immunocompromised patients and in some animal hosts, a higher evolutionary rate and immune-escape related mutations in Spike can be observed. More data of SARS-CoV-2 in immunocompromised patients and viral genomic surveillance in animal hosts are needed to understand the origins of Omicron lineage, BA.2.86, and other potential lineages in the future.

### Continuous improvement in bioinformatics pipelines, especially variant calling, is required to keep pace with novel viruses

Given the unpredictable emergence and spread of novel viruses and variants, the ability to accurately generate high quality genomes and identify true genetic divergence is essential for determining viral epidemiology and pathogenicity, ensuring assay specificity for early diagnostics, and estimating potential drug resistance or immune escape to control clinical disease. Viruses, especially RNA viruses, exhibit high mutation rates, exceptional variation, and rapidly evolve compared to other organisms. This genomic instability leads to divergent genetic features – including SNPs, structural indels, and recombination events – that can be challenging to identify without careful bioinformatic tool selection and parameterization.

SARS-CoV-2 has significantly diverged from the community accepted reference genome, which was isolated from a human clinical sample in 2019 (Genbank accession MN908947.3). The recent identification of the 12nt insertion highlights the importance of routine benchmarking and continuous improvement of SARS-CoV-2 bioinformatics pipelines. The differences and tradeoffs of short read aligners and variant callers has been extensively reviewed elsewhere^31–34^, including benchmarking studies for detecting SARS-CoV-2 in wastewater ^35^. The accuracy of variant callers in these studies varies depending on the type of genetic variation present and the desired result (precision, lineage-defining mutations, novelty detection, etc). It is widely accepted that there is no one-size-fits all pipeline to catch all varieties of genomic divergence. In particular, variant calling using short reads to identify short structural variants (indels <15 nucleotides long) is an outstanding bioinformatics challenge that warrants attention given important public health sequencing projects will be particularly prone to erroneous behavior of variant callers.

Given that variant calling is highly sensitive to input material, sequencing platform, primer scheme, depth of sequencing, bioinformatics tool, and pipeline parameterization, it is nearly impossible to generate a standard bioinformatics pipeline that will remain robust over time. Instead, our recommendation for labs seeking to identify novel variants in near-real time is to run multiple bioinformatics pipelines in parallel and implement different variant callers or variant calling parameters, compare pipeline outputs, then set alerts for manual inspection when significant mismatches occur. Systematic benchmarking studies for variant callers consistently report higher F1 scores when two or more variant callers are implemented and compared for final variant calls. In some studies, the accuracy of variant discovery depended primarily on the variant caller and not the read aligner^33^. Manual inspection can be performed on a bespoke basis for validation, and a polished genome can be reported. Given the relatively small genomes of SARS-CoV-2, this approach should not be significantly more computationally intensive nor impact turnaround time to results. This recommendation is extendable to genomic surveillance for other viral respiratory pathogens.

## Conclusion

The global emergence of BA.2.86 combined with a divergent mutational profile from any previous SARS-CoV-2 variant suggest that BA.2.86 may become an important, dominant variant globally. In addition, BA.2.86 highlighted challenges associated with bioinformatic pipelines and warrant diligent updates to avoid technical artifacts. Lastly, given the precipitous decline in global genomic surveillance, further and strengthened surveillance is needed to assess BA.2.86 potential mutational profile evolution, epidemiological monitoring for clinical severity, and BA.2.86’s contributions to global variant diversity and spread.

## Acknowledgements

We gratefully acknowledge the CDC Traveler-based Genomic Surveillance program for their critical review of this manuscript, specifically Stephen Bart, Cindy Friedman, Dan Payne, Teresa Smith, and Allison Walker. We also gratefully acknowledge all data contributors, i.e., the Authors and their Originating laboratories responsible for obtaining the specimens, and their Submitting laboratories for generating the genetic sequence and metadata and sharing via the GISAID Initiative, on which this research is based.

## Supplementary Information

### Detailed overview of bioinformatics pipelines

#### Bioinformatics analysis

Four custom in-house pipelines were tested to generate a consensus sequence, assign the BA.2.86 lineage, and confirm the lineage-defining 12nt insertion sequence, GTGTGTCATGCCGCTGTTTAAT. The four pipelines vary primarily in variant calling and assembly methods. They are detailed herein:

- **Pipeline 1 implemented the most strict variant calling parameter settings to identify SNPs and implemented soft clipping of adapter sequences to minimize bioinformatics artifacts**. Pipeline 1 was built in SnakeMake (v5.14)^36^. The pipeline and environment executed analysis as follows: raw sequencing reads were demultiplexed (BCL2fastq v2.20; Illumina), confirmed to pass Q30 ≥ 75% (fastqc v0.11.9)^37^, then adaptors were trimmed (trimmomatic v0.36)^38^. Quality trimmed and filtered reads were aligned to the SARS-CoV-2 reference genome, MN908947.3 (bwa v2.2.1)^39^ and filtered based on reads with mapping quality < 30. Primers were softclip from alignments (bamclipper v1.0.0)^40^. Reference genome coverage was calculated (bedtools genomecov v.2.27.1)^41^, and samples less <= 70% reference converge at >= 10X depth were considered for downstream analysis. Variant calling and annotation was performed (freebayes v1.3.6 --ploidy 1 --min-alternate-fraction 0.5 --min-coverage 10)^42^, and variants were filtered based on quality >100 depth >10 and alternative observations/depth >0.5 (bcftools v1.15.1 INFO/AO / INFO/DP > 0.5 & QUAL > 100 & INFO/DP > 10)^43^, and a consensus genome was generated (vcftools vcf-consensus v0.1.14)^44^. Regions with <10x coverage were masked and replaced with “N” (bedtools genomecov v.2.27.1)^41^. Finally, regions outside of the reportable genome range were trimmed (samtools faidx v1.9)^43^.
- **Pipeline 2 implemented more relaxed variant calling parameter settings and omitted soft clipping**. Pipeline 2 was built in Nextflow (v22.04.5)^45^ and used nf-core/viralrecon^46,47^ as a backbone with custom modifications in parameter settings and a change in tooling for variant calling (freebayes v1.3.6)^42^. The pipeline and environment executed analysis as follows: raw sequencing reads were demultiplexed (BCL2fastq; Illumina), confirmed to pass Q30 ≥ 75% (fastqc v0.11.9)^37^, then adaptors were trimmed (fastp v0.23.2)^48^, and human reads were removed (Kraken2 v2.1.2)^49^. Quality trimmed and filtered reads were aligned to the SARS-CoV-2 reference genome, MN908947.3 (bowtie2 v2.4.4)^50^. Primers were removed (iVar v1.3.1)^51^, alignments were sorted and indexed (samtools v1.15.1)^43^, and duplicate reads were marked to ensure high quality alignment (picard v2.27.4^52^ + samtools v1.15.1^43^). Variant calling and annotation was performed (freebayes v1.3.6^42^, SnpEff v5.0^53^, SnpSift v4.3^54^), and a consensus genome was generated based on quality >100, depth >10, and alternative observations/depth >0.5 (bcftools v1.15.1^43^ INFO/AO / INFO/DP > 0.5 & QUAL > 100 & INFO/DP > 10). The SARS-CoV-2 lineage was determined on the consensus sequence using Pangolin (v4.3.1)^55^. Tool parameter settings were used as default except for fastp (--cut_front --cut_tail --trim_poly_x --cut_mean_quality 30 --qualified_quality_phred 30 --unqualified_percent_limit 40 --length_required 50), freebayes (--ploidy 1 --min-alternate-fraction was tested at 0.1, 0.3, and 0,4, --min-coverage 10), and bcftools (INFO/AO / INFO/DP > 0.5 & QUAL > 100 & INFO/DP > 10).
- **Pipeline 3 substituted a variant caller known to perform well on small (<50nt) indels, DeepVariant**. Pipeline 3 used the same tools, order of operations, and parameter settings as Pipeline 2 *except* DeepVariant (version 1.5.0, model_type=WGS)^56^ was used as the variant caller instead of freebayes.
- **Pipeline 4 performed *de novo* assembly in lieu of reference-based assembly**. Pipeline 4 implemented a combination of de novo assembly and read mapping based approach to generate consensus genomes. Briefly we used QC and trimmed reads similar to Pipeline 2 followed by implementation of SPAdes (v3.15.0)^57^ on metagenomic mode (–meta) and remaining default parameters. After de novo assembly of reads, contigs were mapped to SARS-CoV-2 reference genome, MN908947.3, using minimap2 (v2.2.0)^58^ in Geneious Prime (v.2022.1.1; https://www.geneious.com) to visualize read alignments.

**Supplementary Figure 1.**
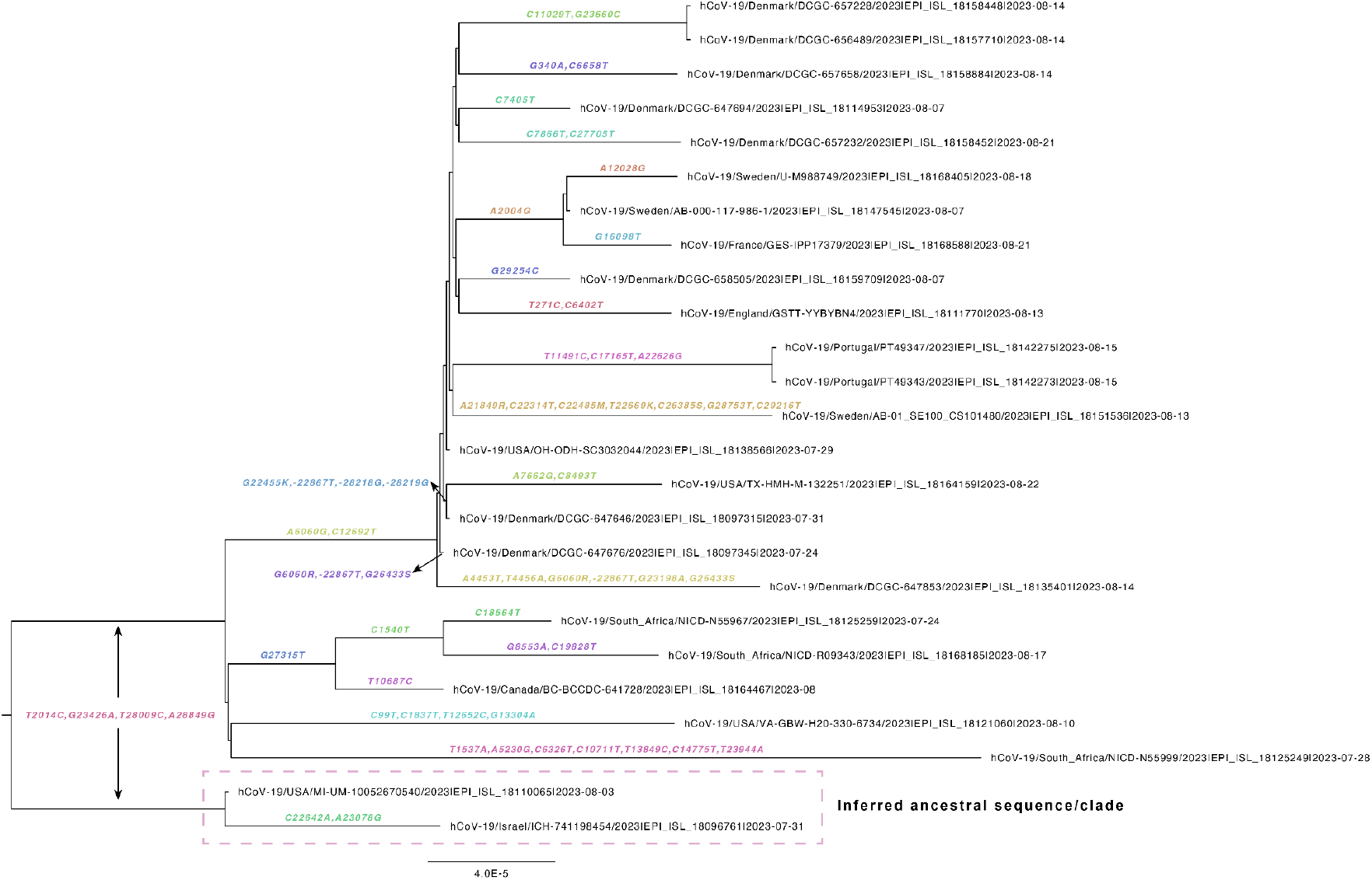
Phylogenetic tree (n=25) with mutations mapped to branches. The inferred ancestral sequence and clade is the USA-MI and Israel isolates, indicating that from evolutionary point of view, these two isolates can be considered as the oldest strain compared to other BA.2.86 sequences in the tree.

